# Human pluripotent reprogramming with CRISPR activators

**DOI:** 10.1101/206144

**Authors:** Jere Weltner, Diego Balboa, Shintaro Katayama, Maxim Bespalov, Kaarel Krjutškov, Eeva-Mari Jouhilahti, Ras Trokovic, Juha Kere, Timo Otonkoski

## Abstract

CRISPR/Cas9 based gene activation (CRISPRa) is an attractive tool for cellular reprogramming applications due to its high multiplexing capacity and direct targeting of endogenous loci. Here we present the reprogramming of primary human skin fibroblasts into induced pluripotent stem cells (iPSC) using CRISPRa, targeting endogenous *OCT4*, *SOX2*, *KLF4*, *MYC* and *LIN28A* promoters. The basal reprogramming efficiency can be improved by an order of magnitude by additionally targeting a conserved Alu-motif, enriched near genes involved in embryo genome activation (EEA-motif). This effect is mediated in part by more efficient activation of *NANOG* and *REX1*. These data constitute a proof of principle that somatic cells can be reprogrammed into iPSC using only CRISPRa. Furthermore, the results unravel previously uncharacterized involvement of EEA-motif-associated mechanisms in cellular reprogramming.

## Introduction

CRISPRa system relies on sequence specific recruitment of a catalytically inactivated version of Cas9 protein (dCas9) to genomic sequences defined by short guide RNA (gRNA) molecules ^1–3^. The fact that dCas9 effectors can be used to control transcription of targeted endogenous loci makes it useful for mediating cellular reprogramming, which requires silencing and activation of endogenous gene sets for proper cell type conversion. CRISPRa may therefore be beneficial in overcoming reprogramming barriers that limit reprogramming efficiency and contribute to the emergence of partially reprogrammed stable cell populations, often associated with inadequate endogenous gene activation or silencing ^4–6^. Previously, dCas9 effectors have been used to mediate differentiation, transdifferentiation and reprogramming of various mouse and human cell types, but complete pluripotent reprogramming using only CRISPRa has not yet been reported ^7–14^.

In addition to gene activation, dCas9 effector mediated DNA targeting can be used to decipher the functions of genomic regulatory elements ^15–17^. Combining reprogramming factor promoter targeting gRNAs with targeting of other regulatory elements has high potential in mediating comprehensive resetting of gene regulatory networks. A conserved Alu-motif was recently reported to be enriched in the promoter areas of the first genes expressed during human embryo genome activation (EGA) ^18^. This sequence is thus likely to be involved in the control of early embryonic transcriptional networks. As human embryos can reprogram somatic cell nuclei^19^, we hypothesized that targeting this EGA enriched Alu-motif (EEA-motif) could enhance reprogramming of somatic cells to pluripotency.

Development of reprogramming approaches for faithful recapitulation of cellular phenotypes is an important task, considering the increasing pace with which reprogrammed cells are moving towards clinical trials ^20^. Here we describe a novel method for reprogramming human cells, including primary adult human skin fibroblasts, into induced pluripotent stem cells by CRISPRa. This opens up important possibilities for the development of more extensive CRISPRa reprogramming approaches for human cells. Efficiency of the method depends on the targeting of the EEA-motif, which results in improved activation of a subset of endogenous genes that work as reprogramming factors, including *NANOG* and *ZFP42* (*REX1*). These data also exemplify the potential in targeting cell type enriched regulatory elements for controlling cell fate.

## Results

### CRISPRa-mediated reprogramming of NSC and EEA-motif targeting

We began human cell reprogramming with CRISPRa using a simplified reprogramming scheme. SpdCas9VP192 mediated *POU5F1* (*OCT4*) activation has been used to replace transgenic OCT4 in human fibroblast reprogramming, while the transgenic expression of only OCT4 has been shown to be sufficient for the reprogramming of neural stem cells (NSC) into iPSC ^12,21^. We therefore combined CRISPRa-mediated *OCT4* activation and NSC reprogramming as an initial model. Since robust CRISPRa reprogramming may depend on high efficiency activation of targeted genes, we built three new trimethoprim (TMP)-inducible dCas9 (DDdCas9) activator constructs containing P65-HSF1 and P300 core domain fusions in addition to the VP192 activator domain ^15,22^ (Fig. 1a). *OCT4* activation in HEK293 cells resulted in heterogeneous OCT4 expression with the activator constructs containing the P300 core (Fig. 1b). Therefore, TetON-DDdCas9VPH was used for *OCT4* activation in NSC reprogramming experiments (Fig. 1c). Addition of doxycycline (DOX) and TMP to iPSC-derived NSCs resulted in the emergence of pluripotent cells, which could be expanded into stable cell lines (Fig. 1d). This demonstrated that CRISPRa mediated activation of endogenous *OCT4* alone was sufficient to reprogram NSC to iPSC.

**Figure 1.**
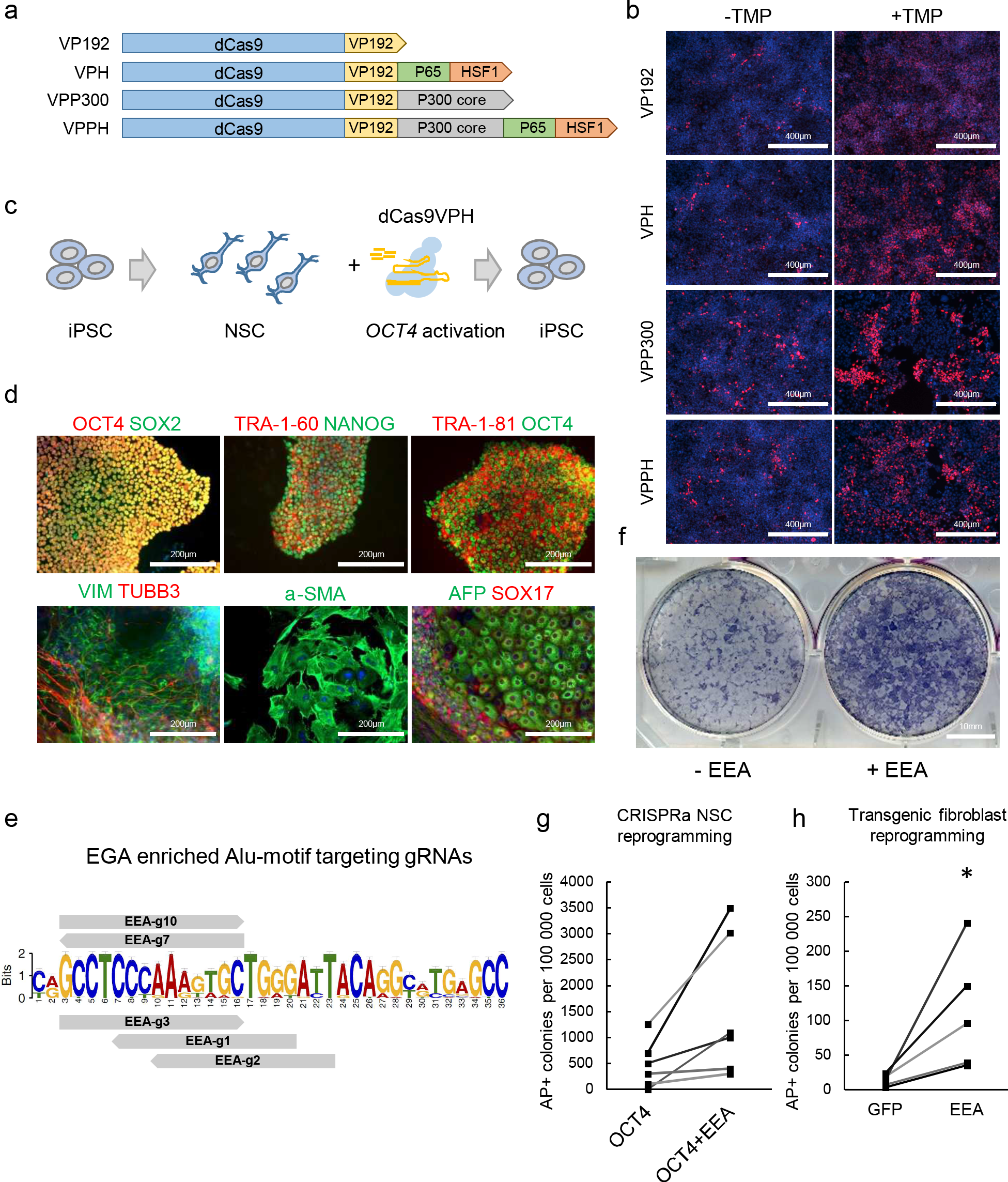
CRISPRa mediated reprogramming of NSCs and EEA-motif targeting. (a) Different dCas9 activator constructs tested. (b) Staining for OCT4 activation with DDdCas9 activators in HEK293 after 5 days of TMP treatment. Nuclei stained blue. (c) Schematic representation of NSC reprogramming into iPSC with dCas9VPH mediated *OCT4* activation. (d) Immunocytochemical detection of pluripotency markers in NCS-derived iPSCs (top row) and tri-lineage differentiation in plated embryoid bodies (bottom row). Nuclei stained blue. (e) Targeting of EGA enriched Alu-motif with SpdCas9 gRNAs. (f) Alkaline phosphatase positive cells induced from NSC by dCas9VPH mediated *OCT4* activation with and without EEA-motif targeting gRNAs. (g) Quantification of iPSC-like alkaline phosphatase positive colonies induced from NSC. n = 6 independent inductions (P=0.053). Data points connected with lines are from the same batch of NSCs. (h) Quantification of iPSC-like alkaline phosphatase positive colonies induced from skin fibroblasts by transgenic reprogramming with GFP control plasmid or EEA-motif targeting gRNA plasmid. n = 5 independent inductions (P=0.034). Data presented as mean ± SEM, two-tailed Student’s t-test. * P<0.05

To determine if EEA-motif targeting could improve CRISPRa reprogramming of NSCs, we designed a set of five gRNAs targeting the 36 bp EEA consensus sequence (Fig. 1e). Addition of the EEA-motif gRNAs in the reprogramming mixture caused a consistent increase in alkaline phosphatase (AP) positive colonies (P=0.053) (Fig. 1f,g). Additionally, reprogramming efficiency of primary human skin fibroblasts, using conventional transgenic episomal reprogramming approach ^23^, was found to increase 8.7-fold over GFP control when simultaneously targeting the EEA-motif with CRISPRa (P=0.034) (Fig. 1h). This suggested that EEA-motif targeting could be useful for improving reprogramming efficiency.

### Fibroblast reprogramming with CRISPRa

To devise a reprogramming system for fibroblasts based solely on CRISPRa, we optimized the promoter targeting of single gRNAs to the canonical reprogramming factors *OCT4*, *SOX2*, *KLF4*, *MYC*, *LIN28A* and *NANOG* in HEK293 ^24,25^ (Fig. 2a,b). Best performing gRNAs targeting *OCT4*, *SOX2*, *KLF4*, *MYC* and *LIN28A* (OMKSL) promoters were concatenated into a single plasmid and tested in selected HEK293 with TMP inducible DDdCas9 activators (Fig. 2c,d). The different activator constructs showed comparable gene activation potential between the targeted reprogramming factors (Fig. 2d).

**Figure 2.**
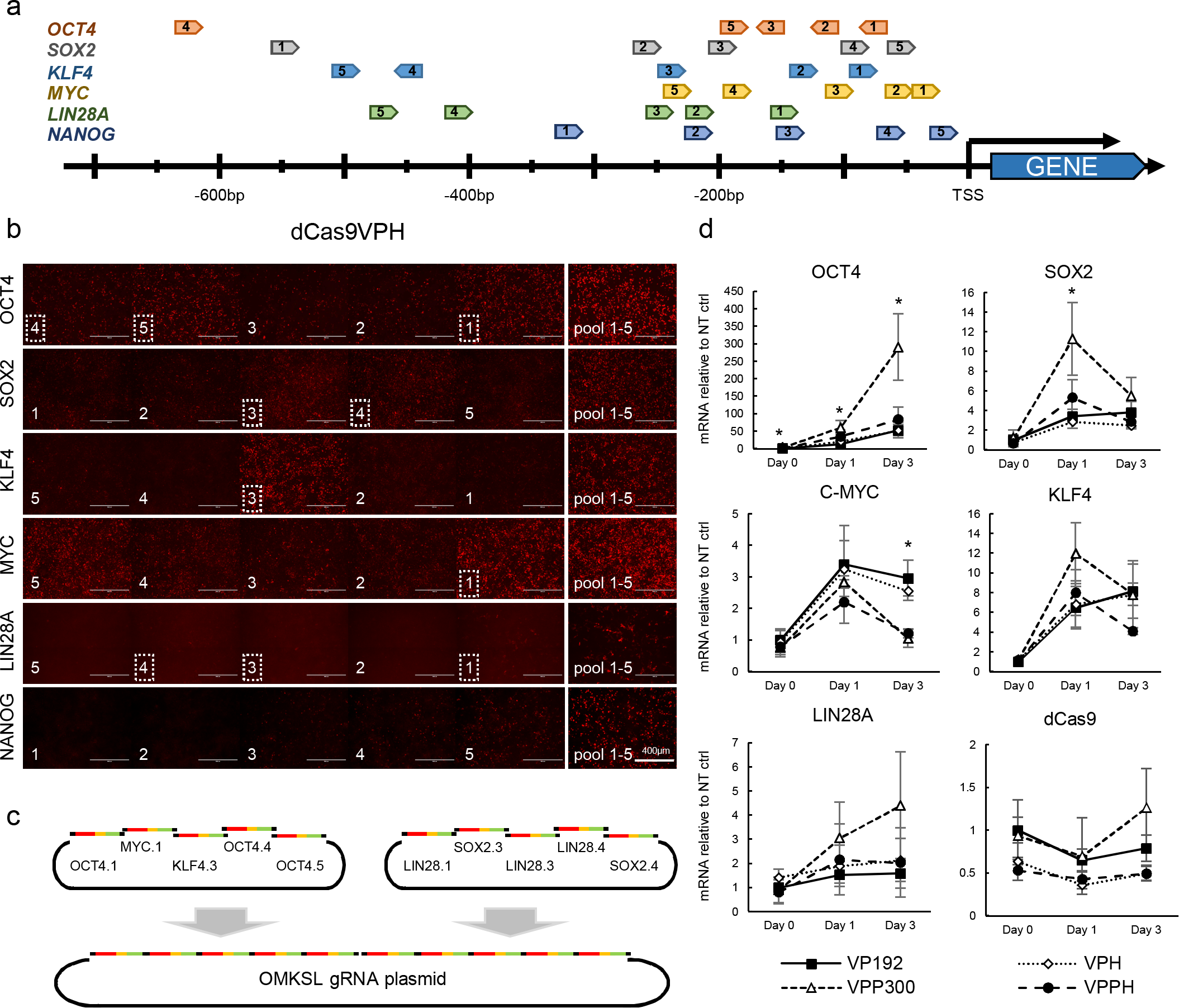
Optimization of dCas9 activator and gRNA targeting in HEK293 for reprogramming factor activation. (a) Locations of promoter targeting gRNAs for reprogramming factors (*OCT4*, *SOX2*, *KLF4*, *C-MYC*, *LIN28A* and *NANOG*) in relation to transcription start site. (b) Immunocytochemical staining of reprogramming factors after single gRNA activation and pooled mixture of five guides in HEK293 with dCas9VPH. Pictures are in similar order to guides in Fig. 2a. Best performing guides used for plasmid cloning are marked with dotted lines. (c) Schematic representation of concatenated reprogramming factor gRNA plasmid construction. (d) Reprogramming factor activation, using constitutively expressed DDdCas9 effectors with different activation domains, in HEK293 by qPCR after TMP addition. n = 3, data are from 3 independent experiments. Error bars represent standard deviation. One way Anova with Tukey HSD test used for statistical comparisons. * P<0.05

Electroporation of primary skin fibroblasts with episomally replicating dCas9VPH plasmid, containing *TP53* targeting shRNA, EEA-motif targeting gRNA plasmid and reprogramming factor targeting gRNA plasmid resulted in the emergence of iPSC-like colonies after 19 days (Fig. 3a see also cell line derivation in the methods). Reprogramming of 72-year-old donor skin fibroblasts was dependent on an additional KLF4 and MYC gRNA encoding plasmid (KM), possibly due to insufficient activation by the single guides used to target these factors in the OMKSL concatenate vector (Fig. 2c see also Supplemental Fig. 3c). The resulting colonies could be expanded into iPSC lines demonstrating typical pluripotency markers and differentiation into three germ layer derivatives *in vitro* and *in vivo* (Fig. 3b and Supplemental Fig. 1a,b). These CRISPRa-induced iPSC lines presented normal karyotypes (Fig. 3c and Supplemental Fig. 1c), and clustered away from HFFs, together with control Sendai viral iPSCs (HEL46.11) and H9 embryonic stem cells, by transcriptional (Fig. 3d,e) and DNA methylation profiles (Fig. 3f). Overall, this demonstrated that CRISPRa reprogramming can be used to derive fully reprogrammed iPSCs from human skin fibroblasts.

**Figure 3.**
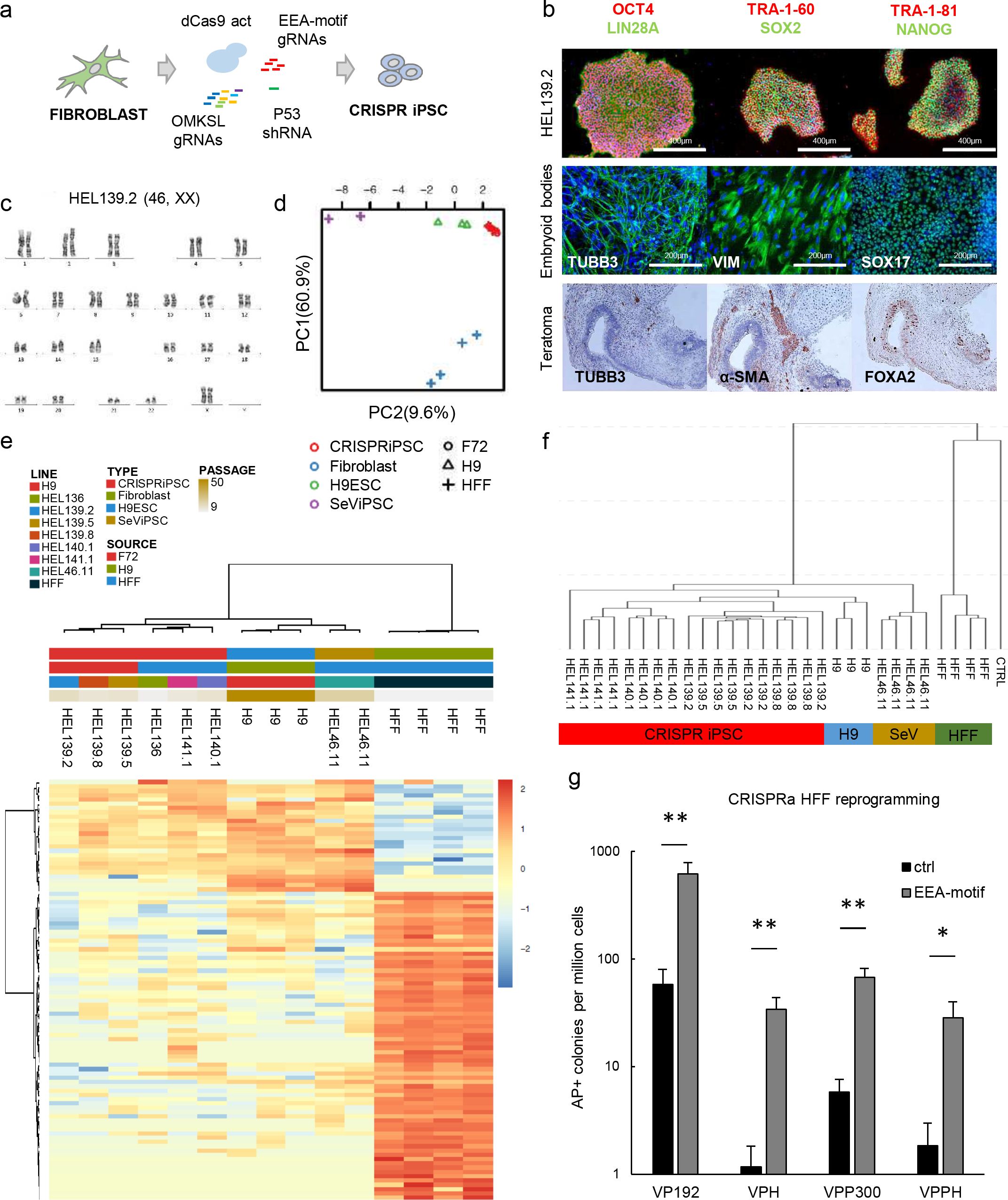
EEA-motif targeting enhances derivation of CRISPRa iPSC from primary skin fibroblasts. (a) Schematic representation of skin fibroblast reprogramming with dCas9 activators. (b) Pluripotency factor expression in CRISPR-iPSC colonies (top row) and tri-lineage differentiation markers for ectoderm (TUBB3), mesoderm (Vimentin and a-SMA) and endoderm (SOX17 and FOXA2) in embryoid bodies (middle row) and teratomas (bottom row). (c) Normal 46, XX karyotype of a CRISPRa iPSC line HEL139.2. (d) Principal component analysis of CRISPR iPSC lines, control PSC lines and HFFs based on expression of 268 significantly fluctuated genes. (e) Clustering of iPSC lines and HFFs based on expression of 165 significantly fluctuated and differentially regulated genes. (f) Clustering of CRISPR iPSC lines and control pluripotent stem cells based on DNA methylation. (g) Effect of different activation domains and EEA-motif targeting on CRISPRa reprogramming efficiency of HFFs. n = 6 from 3 independent experiments. Data presented as mean ± SEM, two tailed Student’s t-test. * P<0.05, ** P<0.01

In order to further optimize the method, we then tested the effect of the different dCas9 activator constructs on reprogramming efficiency using neonatal foreskin fibroblasts (HFF). CRISPRa reprogramming using dCas9VP192 resulted in the most efficient AP positive colony formation (up to 0.062% of electroporated cells) (Fig. 3g). Similar pattern of efficiency was also seen when targeting only *OCT4* for activation in an otherwise transgenic reprogramming approach (Supplemental Fig. 2a). There was an additional negative effect on reprogramming efficiency by the dCas9VPH activator even when no gRNAs were present (Supplemental Fig. 2b). As dCas9VP192 appeared to perform best in the CRISPRa reprogramming of human fibroblasts, it was used in the subsequent experiments.

### Transcriptional analysis of CRISPRa reprogramming

EEA-motif targeting greatly enhanced the CRISPRa reprogramming efficiency, ranging from 10.5-fold increase with VP192 (P=0.009) to 29.2-fold increase with VPH (P=0.007) (Fig. 3g). To decipher the mechanism behind the increase in reprogramming efficiency mediated by the EEA-motif targeting, we conducted expression profile analysis of HFF cell populations undergoing CRISPRa-induced reprogramming in the presence and absence of the EEA-motif gRNAs (Fig. 4a). Based on fluctuated genes in the full data set, the samples clustered primarily by induction date (Fig. 4b). Additionally, by day 12 the OMKSL+KM+EEA gRNA treated cells clustered separately from the rest of the samples and demonstrated higher expression of 78 genes primarily associated with pluripotency and TGF-b signalling (Group 6 in Fig. 4b,c).

**Figure 4.**
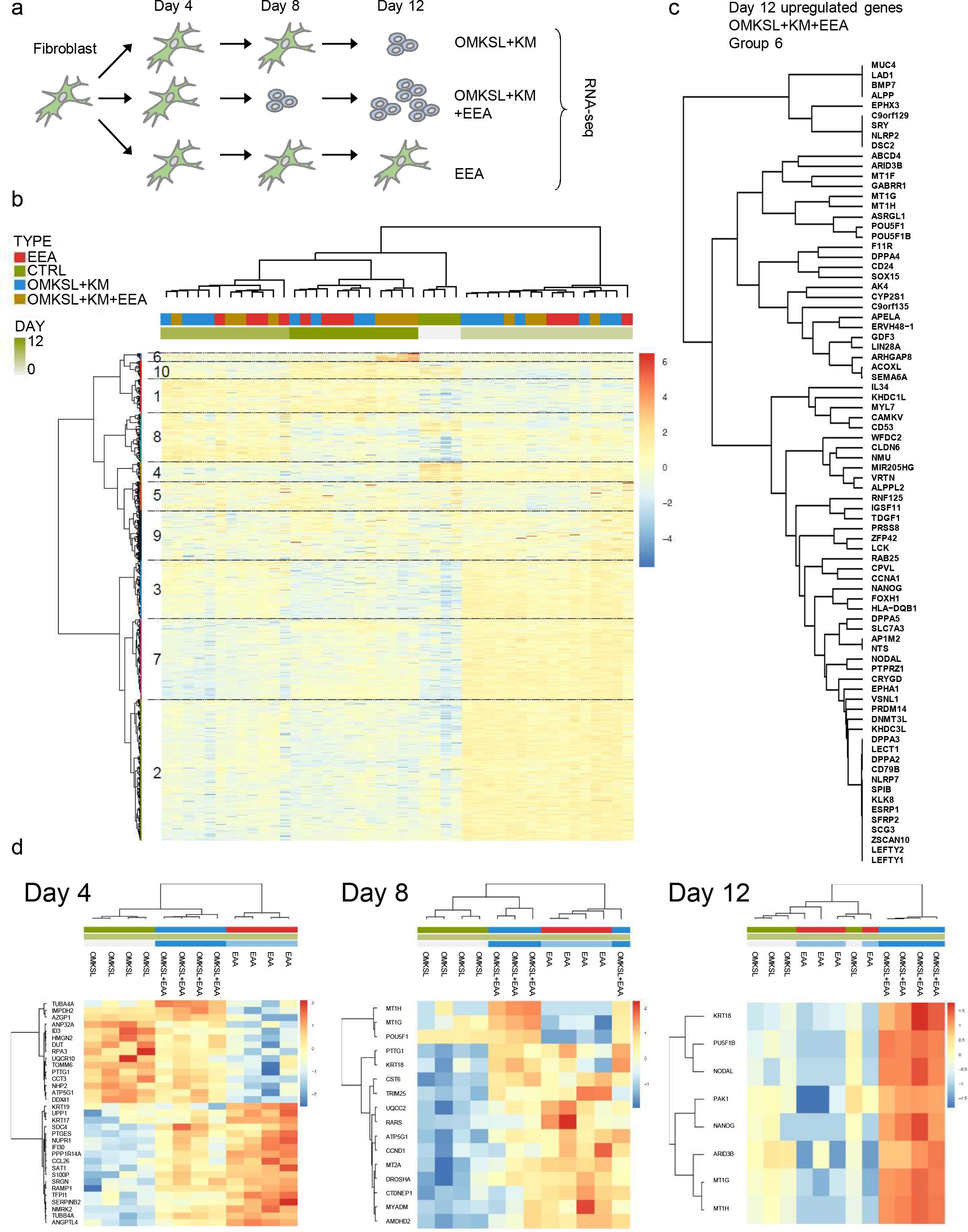
Transcriptional analysis of reprogramming cell populations. (a) Schematic representation of skin fibroblast reprograming with dCas9VP192 for RNA sequencing samples. (b) Clustering of all reprogramming samples based on expression of 4,972 significantly fluctuated genes. Group 6 represents genes upregulated specifically by EEA-guides at day 12. (c) Upregulated genes at day 12 of reprogramming in OMKSL+KM+EEA targeted cells (Group 6). (d) Clustering of samples within day 4, 8 and 12 time points based on differentially regulated genes.

The bulk RNA samples represent heterogeneous cell populations where the majority of the cells are not undergoing complete reprogramming. The clustering seen by induction day may thus reflect nonspecific responses of the fibroblasts to handling, e.g., electroporation. Therefore, we also compared the samples within each time point. This revealed a common set of upregulated genes on day 4 in the conditions containing EEA-motif targeting gRNAs (OMKSL+KM+EEA and EEA, Fig. 4d). These EEA-associated genes had a significantly higher number of EEA-gRNA 1 (EEA-g1) binding sites near their upstream regions (-10kb to +1kb from TSS) (mean 0.409 per kb, n=18, P=5x10^−5^, versus genomic mean 0.215 per kb for protein coding genes) (see also Fig. 6a for EEA-g1). This suggested a preferential initial activation of genes with multiple EEA gRNA target sites. A set of EEA-associated genes was also seen upregulated in the day 8 samples (Fig. 4d), but these genes did not show enrichment for EEA-g1 sites (mean 0.280 per kb, n=13, P=0.068). Significant enrichment was also not detected in the genes that were upregulated by OMKSL+KM only on day 4 (mean 0.224 per kb, n=15, P=0.314) (Fig. 4d), or in the pluripotency-associated genes that were upregulated on day 12 (Group 6, mean 0.248, n=78, P=0.070) (Fig. 4c). However, the day 8 upregulated EEA-associated genes included factors like *TRIM25*, linked to LIN28A function ^26^, and *DROSHA* and *CCND1* which have been associated with cellular reprogramming ^27,28^ (Fig. 4d). EEA-motif targeting at the mid stages of induction may thus contribute to more efficient expression of factors that can promote reprogramming, even if enrichment of EEA-g1 sites was not detected. Unlike day 4 and day 8 upregulated genes, day 12 genes did not show division between EEA related and OMKSL+KM related sets (Fig. 4d), suggesting that the EEA-motif targeting primarily affects the initial stages of the reprogramming process prior to colony formation.

### EEA-associated pluripotent reprogramming factors

Detection of transcriptional changes occurring in small subsets of reprogramming cells can be challenging using RNA-seq of bulk mRNA. This was evident from the absence of detectable LIN28A reads from some of the sequencing samples, although LIN28A protein could clearly be detected by immunostaining in the forming colonies (Fig. 5a). This could also lead to poor detection of reprogramming factors targeted by EEA gRNAs. Assuming that the EEA associated reprogramming factors stay expressed in pluripotent cells, they should be detected more reliably in the day 12 samples, as the fibroblast background is diminished due to expansion of the reprogramming colonies (Fig. 5a). We therefore chose to test a set of seven pluripotency associated factors upregulated by day 12 in the OMKSL+KM+EEA reprogramming data set (Group 6) for their ability to enhance the reprogramming efficiency (Fig. 4c and Fig.5 b). Transgenic expression of NANOG and REX1 in CRISPRa reprogramming in the absence of EEA-motif gRNAs, using optimized reprogramming factor gRNA plasmid (Supplemental Fig. 3), resulted in improved reprogramming efficiency (Fig. 5b). This indicated that NANOG and REX1 could be mediating the EEA-motif targeting effect. Assuming that these factors are downstream effectors of EEA-motif targeting, their direct activation should also be enhanced by EEA-motif gRNAs. Accordingly, dCas9VP192 mediated activation of both *NANOG* and *REX1* promoters in HEK293 resulted in higher expression of these genes in the presence of the EEA-motif gRNAs compared to a TdTomato targeting control gRNA (P=0.009 for *NANOG* and P=0.004 for *REX1)* (Fig. 5c). Both *NANOG* and *REX1* loci contain EEA-g1 binding sites near the genes (Supplemental Fig. 4a,b). *REX1* expression was improved by EEA-gRNAs even when *REX1* gRNAs were replaced with *NANOG* gRNAs, whereas *NANOG* expression was not improved by EEA-gRNAs in the presence of *REX1* gRNAs (Supplemental Fig. 4c). Therefore *REX1* activation by targeting of the EEA-gRNA site near its promoter may represent a direct activation effect, possibly aided by NANOG mediated targeting of *REX1*. On the other hand, *NANOG* activation may be more dependent on additional reprogramming factors or *NANOG* promoter targeting by dCas9 activators. Both *NANOG* and *REX1* thus appear to be downstream targets of the EEA-motif gRNAs, contributing to its effect on improving reprogramming efficiency. Consistent with this, NANOG expression could be detected two days earlier in reprogramming colonies in the presence of the EEA-motif targeting gRNAs (Fig. 5d).

**Figure 5.**
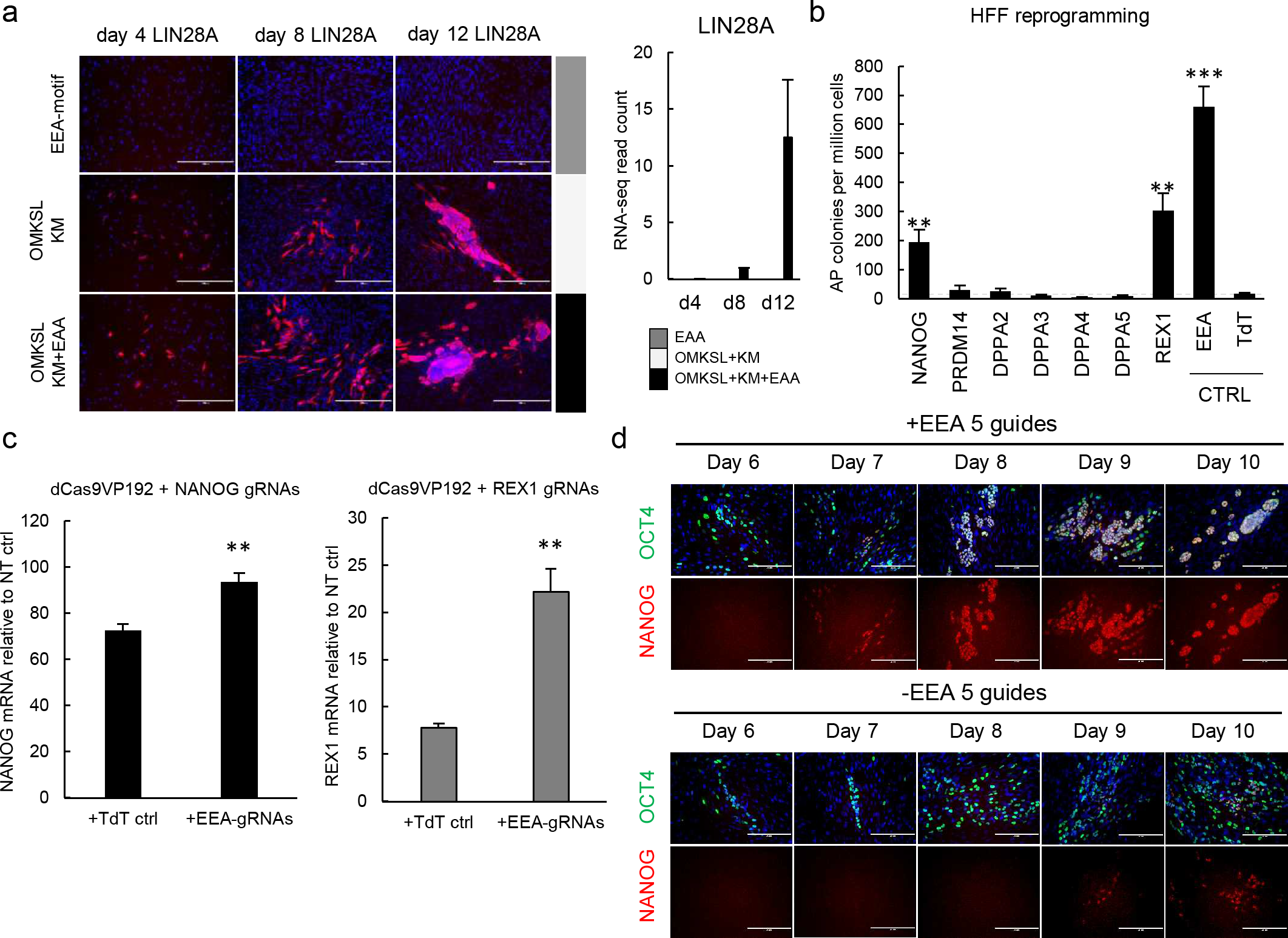
EEA-motif targeting improves *NANOG* and *REX1* activation. (a) Immunostaining of emerging iPSC colonies detects activation of targeted LIN28A before its mRNA reads become detectable in bulk RNA-seq. (b) HFF reprogramming efficiency after transgenic expression of additional factors belonging to Group 6 (Fig. 4c). n=5, 3 independent experiments. (c) qRT-PCR quantification of *NANOG* and *REX1* activation using dCas9VP192 with and without EEA-motif gRNAs in HEK293. n=3, 3 independent experiments. (d) NANOG activation can be detected by immunostaining in CRISPRa reprogramming colonies two days earlier in the presence of EEA-motif targeting guides. Data presented as mean ± SEM, two tailed Student’s t-test. * P<0.05, ** P<0.01

We additionally tested the rest of the reprogramming factors (*OCT4*, *SOX2*, *KLF4*, *MYC* and *LIN28A*), and a set of non-pluripotency associated genes to determine if their activation is affected by simultaneous EEA-motif targeting in HEK293. However, this did not result in consistent effect on transcriptional activation (Supplemental Fig. 4d,e). The EEA-motif targeting thus appears specific only to a subset of reprogramming factors.

### Mechanism of EEA-motif targeting

To further dissect the mechanism behind the EEA-motif targeting, we individually tested all of the five guides targeting the EEA-motif in CRISPRa reprogramming. We also included control guides targeting common guide sequences found in human pluripotent stem cell super enhancers ^29^, to rule out nonspecific global DNA targeting. Of note, the most common gRNAs in these areas also contained multiple Alu sequences and the control guides 8, 9 and 10 also targeted parts of the EEA-motif. Reprogramming efficiency was mainly dependent on the EEA-motif guide 1 (Fig. 6a). There was a noticeable difference in reprogramming efficiencies between EEA-g1 and EEA-g2, which are located next to each other in the EEA-motif consensus (Fig. 1e). These differences may be explained by guide nucleotide composition, as EEA-g2 contained multiple PAM proximal nucleotides that have been shown to be disfavoured ^30^. In accordance with this, EEA-g1 activated a reporter construct with the EEA-motif consensus sequence with higher efficiency than EEA-g2 (P=0.02) (Fig. 6b,c) ^31^. The gRNA nucleotide sequence affecting its efficiency may thus be a crucial determinant in the EEA-motif targeting effect.

**Figure 6.**
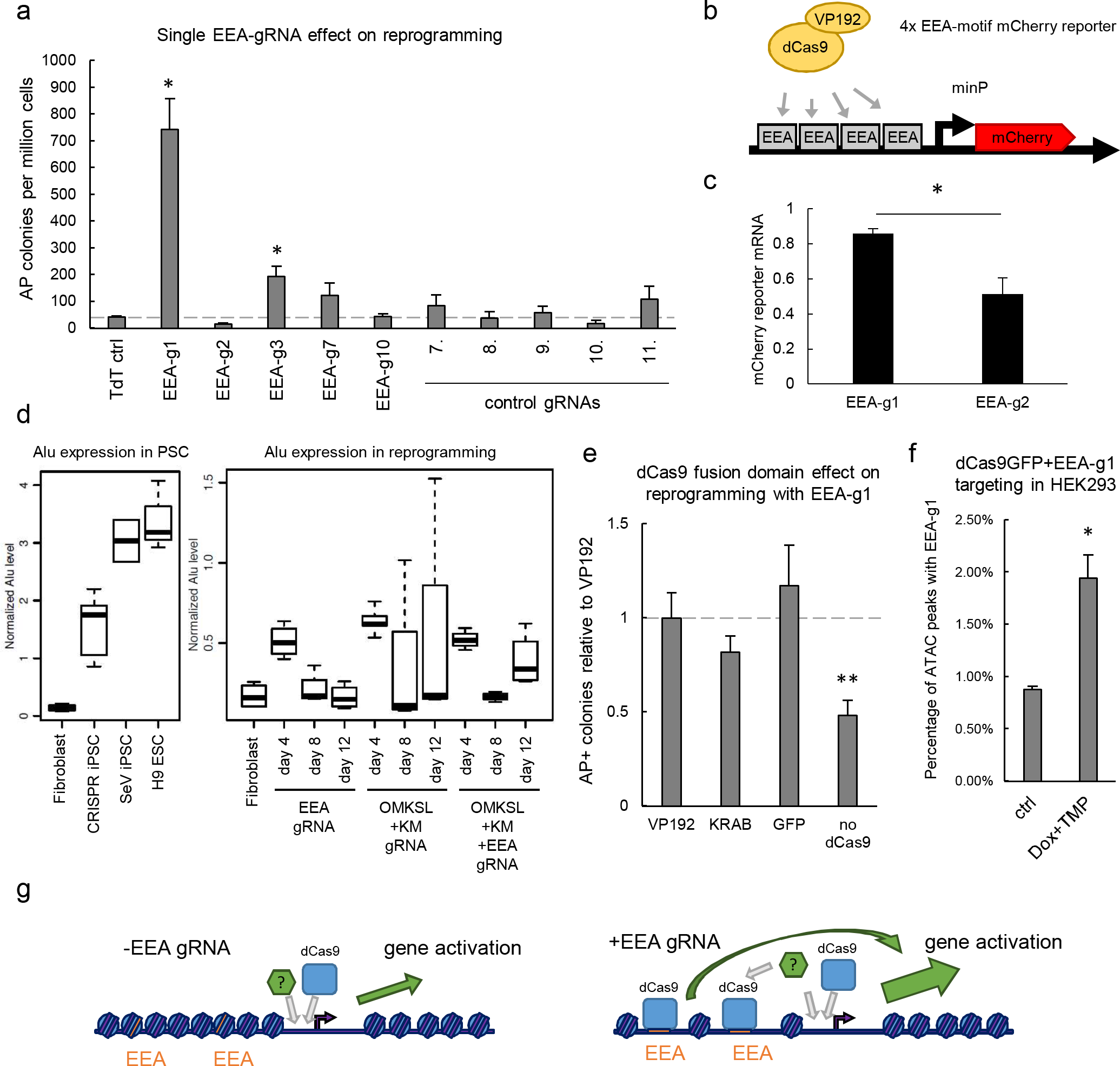
EGA enriched Alu-motif targeting is associated with chromatin opening. (a) Effect of single EEA-motif targeting gRNAs (Fig. 1e) on CRISPRa reprogramming efficiency of HFFs. n=3, data are from 3 independent experiments. (b) Schematic representation of EEA-motif reporter plasmid activation. (c) qRT-PCR quantification of mCherry reporter activation with EEA-motif gRNAs 1 and 2 in reporter transfected HEK293 relative to 5 gRNAs. EEA-g1 activates the reporter with higher efficiency than EEA-g2 (P=0.02). n = 3, 3 independent experiments. (d) Expression of Alu derived transcripts in STRT-seq data of pluripotent stem cells and CRISPRa reprogramming cell populations. (e) Effect of dCas9 fusion domain on reprogramming efficiency in transgenic reprogramming with EEA-g1. n = 6, data are from 3 independent experiments. (f) Increase in the presence of EEA-g1 sequence in ATAC-seq peaks in TetON-DDdCas9GFP and EEA-g1 expressing HEK293. ctrl n=3, DOX+TMP n=5. (g) Schematic model of EEA-motif targeting in gene activation and reprogramming. Data presented as mean ± SEM, two tailed Student’s t-test. * P<0.05, ** P<0.01.

As the EEA-motif is located in the left arm of the Alu consensus sequence, we also looked into expression of Alu sequences that could be detected in the STRT RNA-seq data in the pluripotent stem cells and the reprogramming samples. Alu expression was higher in pluripotent cells than in fibroblasts (Fig 6d). In reprogramming samples Alu expression peaked initially in all day 4 samples, possibly as a response to electroporation, and thereafter decreased (Fig. 6d). Alu expression rose again in the day 12 samples of the OMKSL+KM+EEA samples with higher numbers of pluripotent cell colonies. Overall, EEA-motif targeting did not appear to affect Alu expression in the reprogramming cell batches. Therefore, Alu expression itself is unlikely to explain the potentiating effect of EEA-motif targeting on reprogramming.

We next tested the impact of different dCas9–fused effector domains on the EEA-g1 effect in conventional transgenic reprogramming. Interestingly, the effector domain did not have a significant impact on the reprogramming efficiency, whereas absence of the dCas9 protein resulted in reduced reprogramming efficiency compared to the dCas9VP192 control (P=0.007) (Fig. 6e). The effect of the EEA-motif targeting therefore appears to be mediated specifically by dCas9. It is possible, that dCas9 binding to high affinity guide sites in the EEA-motif may disrupt the chromatin locally to mediate more efficient activation of adjacent genes. As dCas9 binding to DNA has been demonstrated to open chromatin near its target site ^32^, we performed ATAC-seq on samples of HEK293 cells expressing the EEA-g1 and a DOX- and TMP-inducible version of the DDdCas9GFP protein. DOX and TMP treated samples were found to have increased percentage of peaks with overlapping EEA-g1 sites (0.8% in controls, n=3, vs 1.9% in treated cells, n=5, P=0.011) (Fig 6f). This supports a mechanistic model in which dCas9 binding to high efficiency guide sites in the EEA-motif can lead to interference of the local chromatin structure near these elements, which may then contribute to the reprogramming process, for example by aiding in the activation of pluripotency factors with abundant EEA-motif sequences near them (Fig. 6g).

Finally, we tested if the fibroblast CRISPRa reprogramming system could be transferred into inducible transposon based vectors. To this end, we inserted the DDdCas9VP192 activator, under a DOX-inducible promoter, into a PiggyBac vector and the OSK2M2L1 cassette and five guides for the EEA-motif into another one (Supplemental Fig. 3a and Supplemental Fig. 5a). HFFs electroporated with the PiggyBac vectors formed AP positive colonies, which could be expanded into stable iPSC lines, differentiated into three embryonic germ layer derivatives, and re-induced upon DOX and TMP addition (Supplemental Fig. 5). This demonstrated the applicability of the CRISPRa reprograming system in primary cells using various plasmid vectors and the option for the establishment of secondary reprogrammable systems based on CRISPRa.

## Discussion

CRISPR activator approaches hold great potential for controlling cellular reprogramming. The high multiplexing capacity of the system allows simultaneous targeting of large numbers of endogenous genes and genomic control elements using only short guide RNA molecules. This type of approach, combined with large scale synthesis of nucleic acids, can enable comprehensive targeting of gene regulatory networks with great precision for controlling cellular fate. However, until now no robust methods have been described for reprogramming cells into pluripotency by CRISPRa.

We present a method for the efficient conversion of primary human fibroblasts into *bona fide* iPSCs based entirely on the transcriptional control of endogenous genes by CRISPR/dCas9. Activation of core reprogramming factor promoters alone was sufficient but inefficient, whereas additional targeting of a common Alu element brought the efficiency close to established reprogramming methods (Supplemental Fig. 3c). The more complex activator domains did not improve reprogramming efficiency, which mirrors previously reported results for gene activation ^33^, and suggests that the benefit of simple additional fused activation domains may be limited.

It has been estimated that 13% of human genome consists of Short Interspersed Nuclear Element (SINE) sequences, including Alu elements ^34^. Accordingly, EEA-g1 sites can be found in more than 360,000 sites in the human genome. Due to the high abundance of EEA-g1 sites, the motif itself would not be expected to mediate a very strong or specific effect. This is also apparent from the inability of the EEA-gRNAs to reprogram cells by themselves. However, enrichment of the motif sequences near genes may end up enhancing the effect of the motif, as seen in the activation of the genes upregulated at day 4 in an EEA-motif dependent manner. Although we did not detect significant enrichment of EEA-g1 sites near the 78 day 12 upregulated pluripotency associated genes (Group 6), Alu family repeats in general have been reported to be enriched near promoters of pluripotent stem cell expressed genes ^35^. In the reprogramming context this may end up biasing the EEA-motif targeting to preferentially affect pluripotency factors. However, in bulk RNA data this effect may be masked by the background of non-reprogramming cells, and therefore more thorough characterization of the EEA-motif targeting effect will require enrichment for the cell populations that undergo successful reprogramming ^36,37^.

The EEA-motif targeting effect was not dependent on the activator domain of the dCas9 effector in transgenic reprogramming. Instead, the effect may be associated with a dCas9 mediated opening of local chromatin. Alternatively, it is possible that dCas9 binding to the EEA-motif interferes with other factors targeting the motif. Alu elements have been linked with insulator function, including those near KRT18, which was upregulated at day 8 by EEA-motif targeting (Fig. 4d) ^38^. Therefore, the opening of these elements, or interference with their function, may contribute to more efficient activation of nearby genes by interfering with chromatin insulation. Further insight into the mechanisms will require characterisation of factors binding to the motif during reprogramming as well as characterization of the function of early embryo factors, which are known to target the motif, e.g., PRD-like totipotent cell homeodomain factors and HNF4a ^31,39–42^.

In conclusion, CRISPRa reprogramming will provide a powerful tool for inducing pluripotent cells. The core method described here can be further improved by targeting known pluripotency genes and regulatory elements, as well as by screening for novel reprogramming factors and elements ^22,43,44^. This will pave way for the development of more comprehensive CRISPRa reprogramming strategies, which in combination with transgenic factors, RNAi, and small molecular compounds, will promote more efficient and specific reprogramming of human cells for future applications.

## Accession codes

Array Express E-MTAB-6185, E-MTAB-6186, (MethylationEPIC array & ATACseq are in progress)

## Acknowledgements

We thank Jonna Saarimäki-Vire, Solja Eurola, Heli Grym, Anni Laitinen, Minnamari Salmela and Yuval Novik for technical assistance with the work. We thank Ingegerd Fransson for sequencing sample preparation and Anastasius Damdimopoulos for help with the ATAC-seq data processing. We also thank Sanna Vuoristo for comments on the manuscript, and The Mutation Analysis Core Facility (MAF) at the Karolinska University Hospital for their support on the DNA methylation analysis and Bioinformatics and Expression Analysis core facility (BEA) for assistance with sequencing.

This work has been supported by the 3i Regeneration project (number 40395/13; a TEKES Large Strategic Research Opening), Jane and Aatos Erkko Foundation, Academy of Finland (No. 297466), Sigrid Jusélius Foundation, Novo Nordisk Foundation, Instrumentarium Science Foundation, the Doctoral Program in Biomedicine at University of Helsinki, and Knut and Alice Wallenberg Foundation (KAW2015.0096). This study was performed on resources provided by SNIC through Uppsala Multidisciplinary Center for Advanced Computational Science (UPPMAX) under project b2014069.

## Author contributions

Conceptualization, J.W. and D.B.; Methodology, J.W., D.B. and M.B.; Software, S.K.; Formal Analysis, J.W. and S.K.; Investigation, J.W., D.B., M.B., K.K., E.M.J.; Writing – Original Draft, J.W.; Writing – Review & Editing, all authors; Supervision, R.T., J.K. and T.O.; Funding Acquisition, J.K. and T.O.

## Materials and Methods

### Ethical consent

The generation of the human induced pluripotent stem cell lines used in this study was approved by the Coordinating Ethics Committee of the Helsinki and Uusimaa Hospital District (Nro 423/13/03/00/08).

### Cell culture

HEK293s, human foreskin fibroblasts (HFF, ATCC line CRL-2429) and adult human dermal fibroblasts were cultured in fibroblast medium (Dulbecco’s modified Eagle’s medium (DMEM; Life Technologies) containing 10% fetal bovine serum (FBS; Life Technologies), 2mM GlutaMAX (Life Technologies), and 100 μg/ml penicillin-streptomycin (Life Technologies)). Human induced pluripotent cells and embryonic stem cells were cultured on Matrigel (BD Biosciences) coated plates in E8 medium (Life Technologies) and split using 5 μM EDTA. Medium was changed every other day. All cells were kept in an incubator at 37°C and 5% CO2.

### Guide RNA design and production

Guide RNAs were designed and assembled as described by Balboa et al. ^1^. Briefly, guide RNA expression cassettes, containing U6 promoter, chimeric single guide RNA and a PolIII terminator were assembled by PCR and concatenated into plasmids using Golden Gate assembly. Concatenated guide sets were cloned into episomal OriP-EBNA1 containing plasmids for reprogramming experiments. A list of guide RNA oligonucleotides is provided in the Supplemental Table 1.

### dCas9 activator plasmid construction

dCas9VPH construct was cloned by adding a P65-HSF1 containing fragment from lenti-MS2-P65-HSF1_Hygro (gift from Feng Zhang, Addgene Plasmid #61426) after the VP192 domain by PCR. dCas9VPP300 was cloned by PCR amplifying the P300 core domain from human cDNA and cloning it after the VP192 domain, as described by Hilton et al.^2^. dCas9VPPH was cloned by adding the P65-HSF1 domain in fusion after the VP192-P300core domain. Activator plasmids were first cloned into CAG-dCas9VP192-T2A-GFP-IRES-Puro backbone and further cloned into pCXLE-dCas9VP192-T2A-EGFP-shP53 (Addgene plasmid #69535) backbone with XhoI and BsrGI. Plasmids used in this study will be made available on Addgene https://www.addgene.org/Timo_Otonkoski/ see also Supplemental Table 2.

### Cell transfection

HEK293 cells were seeded on tissue culture treated 24 well plates one day prior to transfection (10^5^ cells/well). Cells were transfected using 4:1 ratio of FuGENE HD transfection reagent (Promega) in fibroblast culture medium with 500 ng of dCas9 transactivator encoding plasmid and 100 to 200 ng of guide RNA-PCR or 250 ng of dCas9 transactivator encoding plasmid and 250 ng of concatenated guide RNA encoding plasmid. Cells were cultured for 72 h post-transfection, after which samples were collected for qRT-PCR or immunocytochemical staining. HEK293 cells containing the destabilized dCas9 activators and guides were transfected with dCas9 activator, guide RNA and PiggyBac transposase plasmids, 100 ng of each, and selected with Puromycin (2 mg/ml; Sigma) and G418 (5 mg/ml; Life Technologies).

### Quantitative Reverse Transcription Polymerase Chain Reaction

Total RNA was extracted from cells using NucleoSpin Plus RNA kit (Macherey-Nagel). RNA quality and concentration was measured by spectrophotometry using SimpliNano (General Electric). One microgram of total RNA was denatured at 65° C for 1 min and used for reverse transcription (RT) with 0.5 μL Moloney murine leukemia virus (MMLV) reverse transcriptase (M1701, Promega), 0.2 μL Random Primers (C1181, Promega), 1 μL Oligo(dT)18 Primer (SO131, ThermoFIsher) and 0.5 μL Ribolock RNAse inhibitor (EO0382, ThermoFisher) for 90 min at 37° C. For qRT-PCR reactions, 50 ng of retrotranscribed RNA were amplified with 5 μL of forward and reverse primer mix at 2 μM each using 5× HOT FIREPol EvaGreen qPCR Mix Plus (no ROX) in a final volume of 20 μL. QIAgility (Quiagen) liquid handing system was used for pipetting the reactions into 100 well disc that were subsequently sealed and run in Rotor-Gene Q (Qiagen) with a thermal cycle of 95° C for 15 min, followed by 40 cycles of 95° C, 25 s; 57° C, 25 s; 72° C, 25 s, followed by a melting step. Relative quantification of gene expression was analysed using ΔΔCt method, with cyclophilin G (*PPIG*) as endogenous control and an exogenous positive control used as calibrator. Expression levels are relative to non-treated cells or to hESC as indicated in the figure legends. A list of primers used is provided in the Supplemental Table 3.

### NSC differentiation

Human neuroepithelial stem cells (NSC) were derived by differentiating human iPSC HEL24.3 ^3^, and HEL46.11 lines using small molecule cocktail as described elsewhere ^4^, with minor adjustments. iPSCs were detached with StemPro Accutase (Thermo Fisher Scientific) and dissociated gently into single cells suspension in hES-medium in the presence of 5 mM ROCK inhibitor (ROCKi; Y-27632, Selleckchem), 10 mM SB431542 (SB; S1067, Selleckchem), 1 mM dorsomorphin (DM; P5499-5MG, Sigma), 3 mM CHIR-99021 (CHIR; Tocris) and 0,5 mM purmorphamine (PMA; 04-0009, Stemgent) After two days, medium was changed to N2B27 medium (DMEM/F12:Neurobasal (1:1) supplemented with N2 and B27 without vitamin A, NEAA, PenStrep (all Thermo Fisher Scientific) and heparin (2 μg/ml; H3149-50KU, Sigma)) containing the same small molecule cocktail as above. On day 4, SB and DM were withdrawn and 150 μM ascorbic acid (AA) was added to N2B27. On day 6, the neurospheres were dissociated with 1 ml pipette and plated on Matrigel in N2B27 media containing AA, CHIR and PMA (growth media). First two passages were split at 1:3 ratio and cells were plated into growth media containing 5 mM ROCKi, which was removed next day. Later passages were split with 1:10 and 1:20 ratio using StemPro Accutase. Media was changed every other day.

### NSC reprogramming

NSCs were grown for at least five passages before electroporation. For electroporation, cells were detached with StemPro Accutase and dissociated into single cells. Cells were washed once with PBS and electroporated with Neon Transfection system (Invitrogen). Two million cells were used per electroporation using 100 μl tips with 1300 V, 30 ms, one pulse settings. 2 mg of PB-tight-DDdCas9VPH-GFP-IRES-Neo activator plasmid, 1.5 mg PB-GG-OCT4-1-5-PGK-Puro gRNA plasmid and 1.5 mg PB-GG-EEA-5g-PGK-Puro gRNA plasmid were used with 0.5 mg PiggyBac rtTA and 0.5 mg of PiggyBac transposase plasmids. One million electroporated cells were plated per 35mm plate coated with Matrigel in N2B27 media supplemented with 5 mM ROCKi and 10 ng/ml of basic FGF (bFGF, PeproTech). ROCKi was removed the next day. Two days after electroporation cells were treated with Puromycin (0.5 mg/ml; Sigma) and G418 (200 mg/ml; Life Technologies) for 5 days. On day 8 after electroporation reprogramming was initiated by adding doxycycline (DOX, 2 mg/ml; Sigma) and trimethoprim (TMP, 1 mM; Sigma). After 5 days of induction media was changed to hES-medium gradually over a week. During the conversion process cells were split 3 times. On day 18 of induction cells were fixed with 4% paraformaldehyde (PFA) for AP staining or picked for iPSC derivation. Media was changed every other day.

### Fibroblast reprogramming

Human skin fibroblasts were detached as single cells from the culture plates with TrypLE Select (Gibco) and washed with PBS. Cells were electroporated using the Neon transfection system (Invitrogen). 10^6^ cells and 6 μg of plasmid mixture, containing 2 μg of dCas9 activator plasmid and 4 μg of guide plasmids, were electroporated in a 100 μl tip with 1650 V, 10 ms and 3× pulse settings. Electroporated fibroblasts were plated on Matrigel coated 100 mm diameter cell culture plates in fibroblast medium. After 4 days cell culture medium was changed to 1:1 mixture of fibroblast medium and hES-medium (KnockOut DMEM (Gibco) supplemented with 20% KO serum replacement (Gibco), 1% GlutaMAX (Gibco), 0.1 mM beta-mercaptoethanol, 1% nonessential amino acids (Gibco), and 6 ng/ml basic fibroblast growth factor (FGF-2; Sigma)) supplemented with sodium butyrate (0.25 mM; Sigma). When first colonies started to emerge, cell culture medium was changed to hES-medium until colonies were picked. For iPSC line derivation, colonies were picked manually and plated on Matrigel coated wells in E8 medium. Media were changed every other day. For PiggyBac reprogramming, HFF were electroporated as described above with PB-tight-DDdCas9VP192-GFP-IRES-Neo, PB-CAG-rtTA-IRES-Neo, PB-GG-EEA-5g-OSK2M2L1-PGK-Puro and PiggyBac transposase plasmids. Electroporated cells were plated on cell culture dishes in fibroblast medium. Five days after electroporation cells were selected with Puromycin (1 mg/ml; Sigma) and G418 (5 mg/ml; Roche) for two days after which the selection antibiotic amounts were halved. Selected cells were induced as described above in the presence of TMP (1 μM) and DOX (2 μg/ml). Fresh DOX was supplemented daily. For RNA sequencing samples, passage 10 foreskin fibroblasts were electroporated with pCXLE-dCas9VP192-GFP-shP53 and combinations of GG-EBNA-OSKML-PP, GG-EBNA-KM-PP and GG-EBNA-EEA-5guides-PP plasmids. 250k (day 4 samples) to 125k cells (day 8 and 12 samples) were plated on Matrigel coated 6-well culture plates per well and induced as described above.

### Pluripotent cell line derivation

HEL136 was derived from neonatal male skin fibroblasts (HFF) with GG-EBNA-OMKSL-PP and GG-EBNA-EEA-5guides-PGK-Puro plasmids. This cell line was not stable in long term culture. HEL139 clones were derived from adult female skin fibroblasts (F72) using GG-EBNA-OMKSL-PP, EBNA-EEA-5guides-PGK-Puro and an additional GG-EBNA-KM-PP plasmid (KLF4 and MYC five guides each). HEL140 was derived from neonatal male skin fibroblasts (HFF) with GG-EBNA-OMKSL-PP, EBNA-EEA-5guides-PGK-Puro and GG-EBNA-KM-PP plasmids. HEL141 was derived from neonatal male skin fibroblasts (HFF) with GG-EBNA-EEA-5guides-PGK-Puro, GG-EBNA-KM-PP and GG-EBNA-OS-PP plasmids (OCT4 and SOX2 five guides each). The above cell lines were derived using pCXLE-dCas9VPH-T2A-GFP-shP53 activator plasmid. HEL144 was derived from neonatal male skin fibroblasts (HFF) with inducible PiggyBac vectors using PB-tight-DDdCas9VP192-GFP-IRES-Neo activator and PB-EEA-5g-OSK2M2L1-PGK-Puro guides. Control cell line HEL46.11 was derived from HFFs using CytoTune Sendai Reprogramming Kit (Life Technologies) according to manufacturer’s instructions.

### Alkaline phosphatase staining

iPSC colonies were fixed with 4% paraformaldehyde (PFA) solution for 10 min and washed with phosphate buffered saline (PBS). Thereafter cells were stained in NBT/BCIP (Roche) containing buffer (0.1 M Tris HCl pH 9.5, 0.1 M NaCl, 0.05 M MgCl_2_) until precipitate developed. Reaction was stopped by washing the plates with PBS.

### Immunocytochemistry

Cells were fixed with 4% PFA, permeabilized using 0.2% Triton X-100 and treated with Ultra Vision block (ThermoFisher). Primary antibodies were diluted in 0.1% Tween-20 PBS and incubated either overnight at 6°C with the given dilutions or 2 days in 6°C with halved primary antibody amounts. Secondary antibody incubations were done in room temperature for 30 minutes in the presence of Hoechst33342 to stain the nuclei. Primary antibodies used were: LIN28A (1:250, D84C11 and D1A1A, Cell Signaling), NANOG (1:250, D73G4, Cell Signaling), OCT4 (1:500, sc-8628, Santa Cruz), SOX2 (1:250, D6D9, Cell Signaling), KLF4 (1:250, HPA002926, Sigma-Aldrich), C-MYC (1:250, D3N8F, Cell Signaling; 1:250, [Y69] ab32072, Abcam), TRA-1-60 (1:50, MA1-023, ThermoFisher), TRA-1-81 (1:100, MA1-024, ThermoFisher) TUBB3 (1:500, MAB1195, R&D Systems), AFP (1:400, A0008, Dako), SMA (1:200, A2547, Sigma), VIMENTIN (1:500, sc-5565, Santa Cruz). SOX17 (1:500, AF1924, R&D Systems). Secondary antibodies used were: AlexaFluor 488: donkey anti-goat (1:500, A11055 and 11058; Invitrogen), donkey anti-mouse (1:500, A21202 and A21203; Invitrogen) and donkey anti-rabbit (1:500, A21206 and A21207; Invitrogen).

### Embryoid Body assay

iPSCs were split into small clumps and plated on low attachment dishes (Corning) in hESC medium without bFGF to allow embryoid body (EB) formation. The EB culture medium was supplemented overnight with 5 μM ROCK inhibitor (Y-27632, Selleckchem) after the initial cell plating to improve cell viability. Medium was changed every other day. EBs were grown in suspension for 14 days, after which they were plated on gelatin coated cell culture dishes. EBs were allowed to form outgrowths for 7 days after which cells were fixed with 4% PFA for 30 min and permeabilized using 0.2% Triton X100 (Sigma) in PBS for 30 min. Fixed and permeabilized EB outgrowths were stained as described above.

### Teratoma Assay

About 200,000 morphologically intact iPSC at passage 23 were intratesticularly injected into male NMRI nude mice (Scanbur). The resulting tumours were collected 2 months after injection, fixed with 4% PFA, and hematoxylin and eosin stained. Animal care and experiments were approved by the National Animal Experiment Board in Finland (ESAVI/9978/04.10.07/2014).

### DNA methylation assay and analysis

For DNA methylation array samples, passage 11-20 CRISPRa iPSCs, passage 35 SeV iPSCs and passage 50 H9 ESCs were collected, 1 well of 6-well plate each, as well as HFF control fibroblasts. DNA was purified using DNeasy Blood & Tissue Kit (Qiagen) and the concentrations were adjusted to 11 ng/ μl using Qubit assay (Thermo Fisther Scientific). PicoGreen Assay (ThermoFisher) was used for subsequent normalization of samples. DNA of the samples and three controls (Zymo_low, Zymo_high and 1331-1 CEPH) was treated with sodium bisulphite using the EZ DNA methylation kit (Zymo Research). DNA methylation was quantified using the Illumina Infinium HumanMethylationEPIC BeadChip on an Illumina iScan System using the manufacturer’s standard protocol. Raw IDAT files were processed with Illumina’s GenomeStudio v2011.1, and normalized beta-values, except two controls (Zymo_low and Zymo_high), were applied for the clustering of the DNA methylation profile.

### STRT-sequencing

RNA samples were collected in Trizol Reagent (Life Technologies), 100 μl of chloroform per 500 ml of sample was added and mixed with Trizol Reagent. After centrifugation (12 000 g for 15 min) the transparent upper phase was collected and RNA was further purified using NucleoSpin RNA kit (Macherey-Nagel). RIN values were measured by Bioanalyzer (Agilent) and total-RNA concentrations scaled to equal (10 ng/μl) using Qubit assay (Thermo Fisher Scientific). Bulk-RNA transcriptome analysis was performed by the STRT RNA-seq method^5^ with minor modifications^6^. Briefly, ten nanogram of high-quality input RNA was converted to cDNA and amplified to form an Illumina-compatible 46-plex library. In total, 25 PCR cycles were used, but as six base-pair unique molecular identifiers (UMIs) were applied, only the absolute number of unique reads was calculated per analyzed sample. The library was sequenced by three lanes of Illumina HiSeq2000 instrument.

### STRT data analysis

The sequenced raw STRT reads were processed by STRTprep ^6^; v3dev branch, d7efcde commit (https://github.com/shka/STRTprep/tree/v3dev). In brief, redundant reads after demultiplexing were excluded according to UMI, and the nonredundant reads were aligned to hg19 human reference genome sequences, ERCC spike-in sequences and human ribosomal DNA unit (GenBank: U13369). Uniquely mapped reads within (i) the 5’-UTR or the proximal upstream (up to 500 bp) of the RefSeq protein coding genes, (ii) the 5’-UTR or the proximal upstream of some PRD genes, which were not yet defined by RefSeq ^7^, and (iii) within the first 50 bp of spike-in sequences, were counted. The processed reads were aligned also to Alu canonical sequence (http://www.repeatmasker.org/AluSubfamilies/humanAluSubfamilies.html) using the same methodology with STRTprep, to count Alu transcripts.

Significance of fluctuation on gene expression was tested by comparison with fluctuation of spike-in levels as ^6^. Differential expression between the sample types were tested by SAMstrt ^8^. Regulated genes were selected by corrected fluctuation p-value < 0.05 (as significant degree of expression change) and differential expression q-value < 0.05 (as significant contrast between the types), except Fig. 4b. In contrast, genes in Fig. 4b were selected only by the corrected fluctuation p-value < 0.05; this approach might select also genes independent from the samples types (ex. batchs, circadian, cell-cycle etc); however, the unsupervised clustering at the Fig. 4b elucidated that time points in the reprogramming were the major factor relevant to the expression changes. Hierarchical clustering of normalized expression profiles and the illustration were performed by aheatmap function in NMF package ^9^ via heatmap_diffexp plugin of STRTprep; color gradient in the heatmap represents Z-score of each gene, and the profile was clustered by Ward’s algorithm on Spearman correlation based distance. Principal component analysis was performed on fluctuated genes via pca plugin of STRTprep.

### ATAC-sequencing

HEK293 cell lines used for ATAC-seq samples were transfected with PiggyBac vectors for TetON-DDdCas9GFP-IRES-Neo, CAG-rtTA-IRES-Neo and 36bp-guide1-PGK-Puro. Cells were selected with Puromycin and G418, after which cells were sub cloned from single cells and clones were selected based on homogenous GFP expression upon doxycycline addition. For sample preparation, cell clones were treated for three days with doxycycline (1 μg/ml) and trimethoprim (1 μM). Fifty thousand cells were collected for ATAC-seq library preparation according to Buenrostro et al. ^10^. Briefly, nuclei were extracted by continuous centrifugations and chromatin was exposed for enzymatic tagmentation and 12-plex Illumina-compatible library. All fragments were amplified by Phusion hot-start polymerase using 12 cycles of PCR in total. The ATAC-libary was sequenced on one lane of Illumina HiSeq2000 instrument. Peak calling was done with Homer looking for histone like peaks, using sample 5 (non-treated control) as background.

### Statistical Analysis

Statistical analysis was performed as described in the figure legends P-values of less than 0.05 were considered significant (* P<0.05, ** P<0.01, *** P<0.001). Estimating EEA-g1 enrichment near upstream regions of EEA associated genes was done by Monte Carlo sampling with 10^5^ random permutations of n number of genes (as defined in the text) from a pool of 19806 protein coding genes.

---

**Figure S1.**
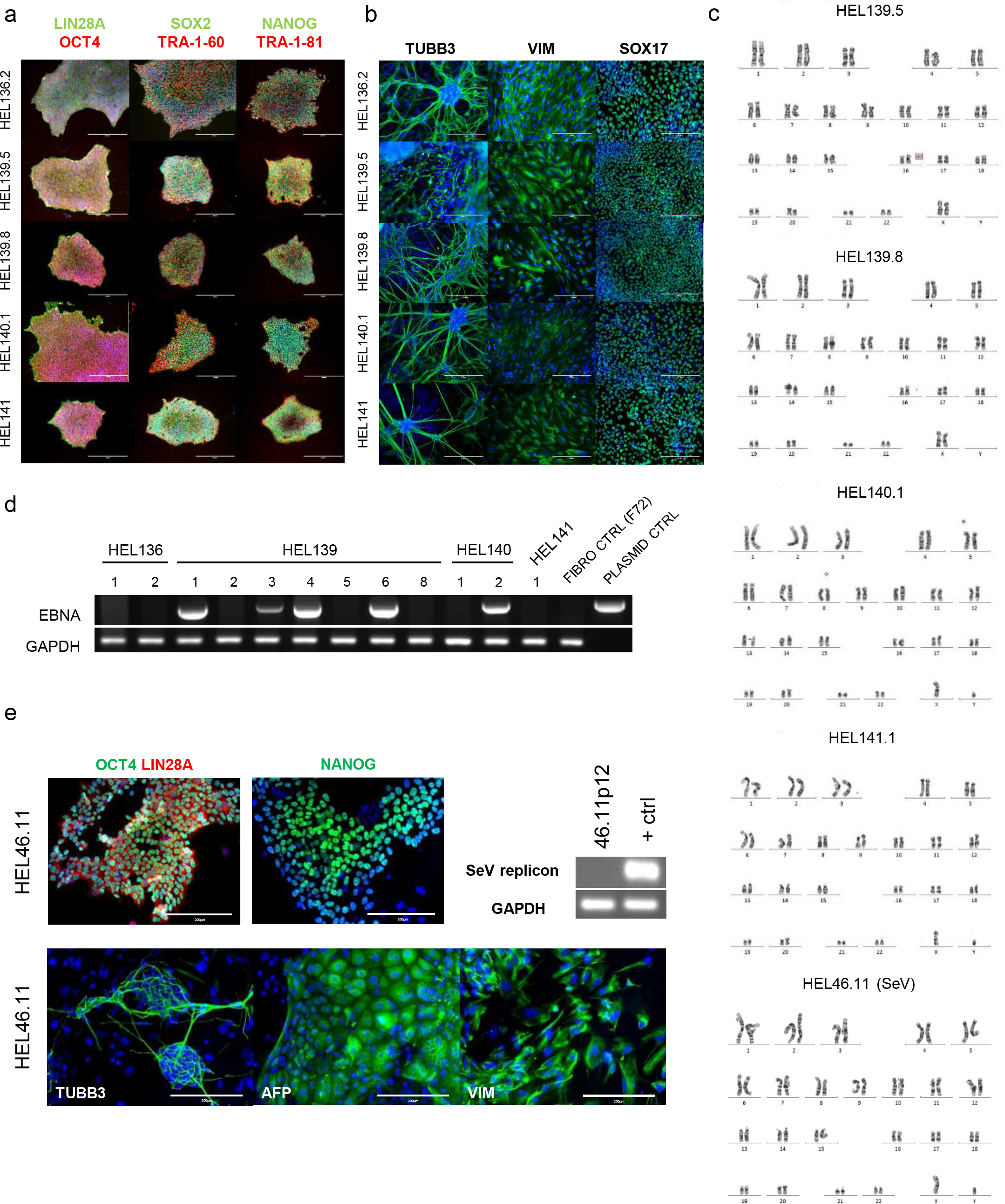
Pluripotency characterization of iPCS lines, (a) Episomal plasmid derived CRISPR iPSCs from HFFs (HEL136, 140, 141) and F72 fibroblasts (HEL139) are positive for pluripotency associated factors. Scale bar 400μm (b) Embryoid body differentiation of the cell lines into three embryonic germ layers: ectoderm (TUBB3), mesoderm (VIMENTIN) and endoderm (SOX17) (green). Scale bar 200μm (c) Karyotypes of HEL139.5 (46, XX), HEL139.8 (46, XX), HEL140.1 (46, XY, 75% of analysed cells), HEL141 (46, XY, abn(3q)) and SeV control line HEL46.11 (46, XX). (d) Episome detection by PCR with EBNA primers of CRISPR dCas9VPH reprogrammed iPSCs at passage 3. (e) Pluripotency, differentiation and absence of viral episome characterization of control SeV iPSC line HEL46.11. Scale bar 200μm. Nuclei stained blue.

**Figure S2.**
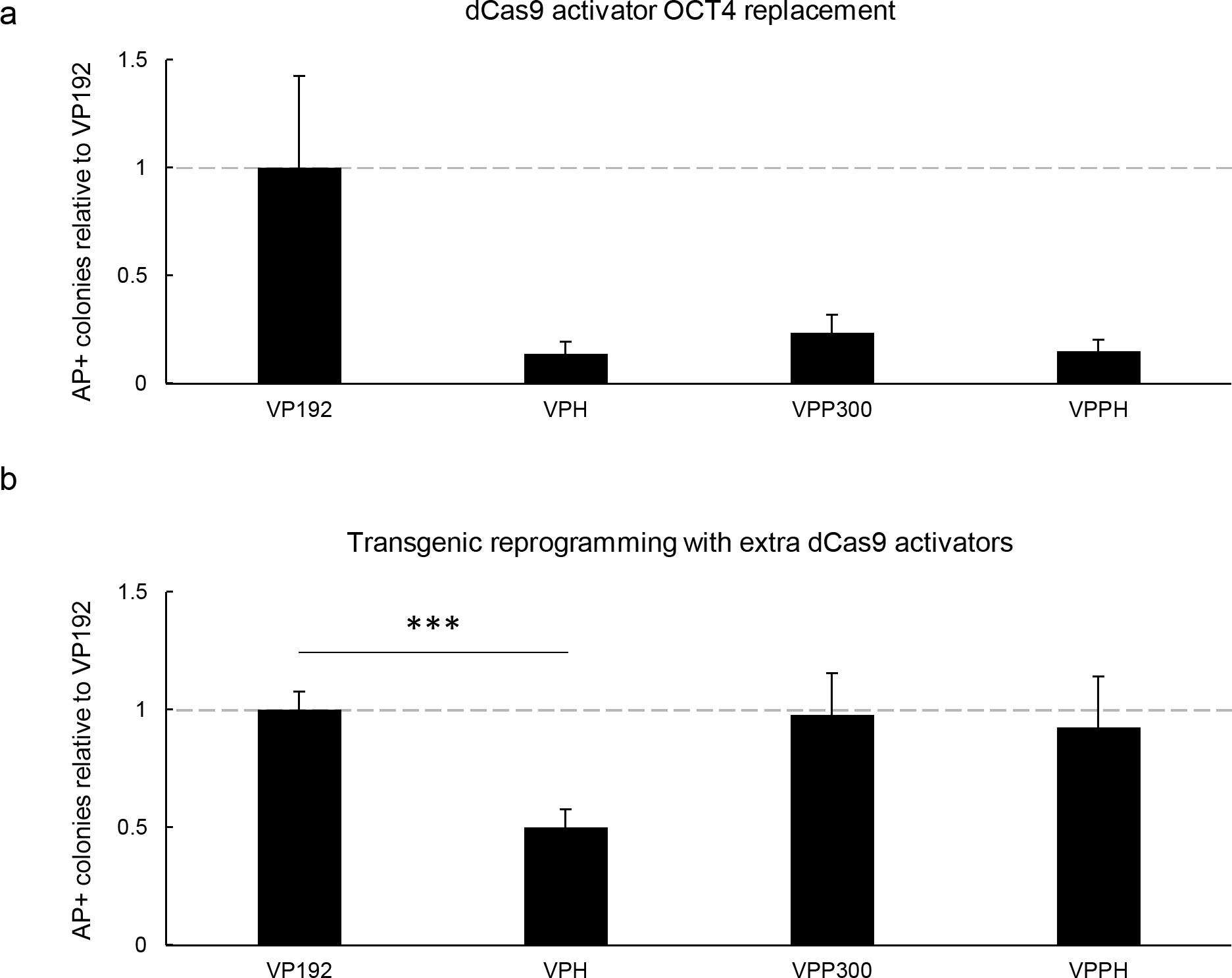
Effect of dCas9 activators on reprogramming efficiency. (a) Reprogramming of HFF with transgenic SOX2, KLF4, L-MYC and LIN28A with dCas9 activator mediated *OCT4* targeting. n = 6, 3 independent experiments. (b) Reprogramming of HFF with transgenic OCT4, KLF4, L-MYC and LIN28A with extra dCas9 activator plasmid without gRNAs. n = 4, 3 independent experiments. Data presented as mean ± SEM, two tailed Student’s t-test. *** P<0.001

**Figure S3.**
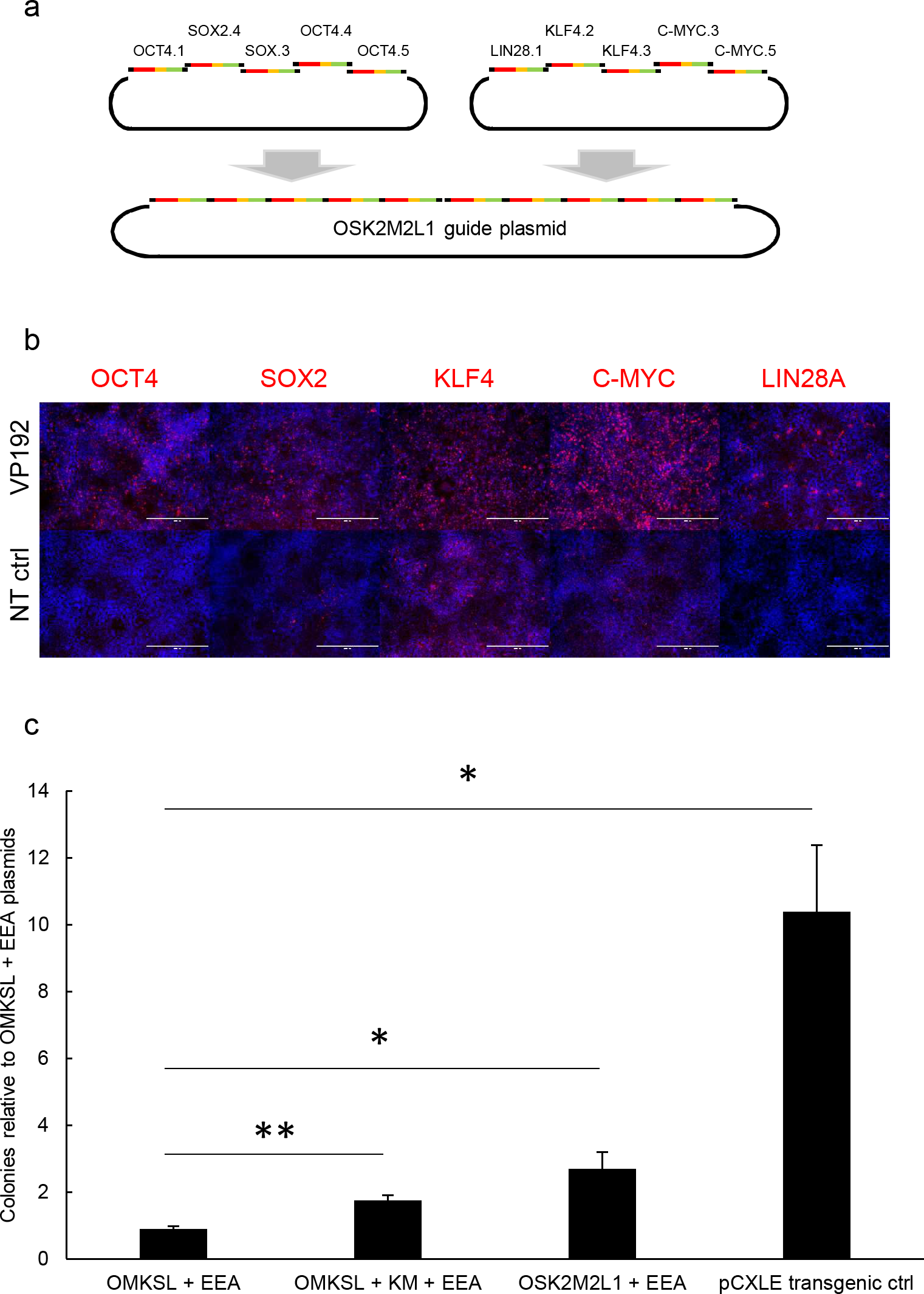
Core reprogramming factor guide optimization for single plasmid delivery. (a) Schematic representation of concatenated core guide plasmid OSK2M2L1 construction. Notice different number of LIN28A, KLF4 and C-MYC guides compared to OMKSL plasmid (Fig. 2) (b) Validation of target gene activation with core factor plasmid in HEK293. Cells transfected with the concatenated reprogramming factor guide plasmid and dCas9–VP192 plasmid, and non-transfected controls (NT). Scale bar 400μm. Nuclei stained blue. (c) Reprogramming efficiencies of HFFs relative to OMKSL and EEA-motif gRNA plasmid. KM plasmid contains five gRNAs targeting KLF4 and five gRNAs targeting MYC. Error bars represent SEM, n = 6, 3 independent experiments. Data presented as mean ± SEM, two tailed Student’s t-test. * P<0.05, ** P<0.01

**Figure S4.**
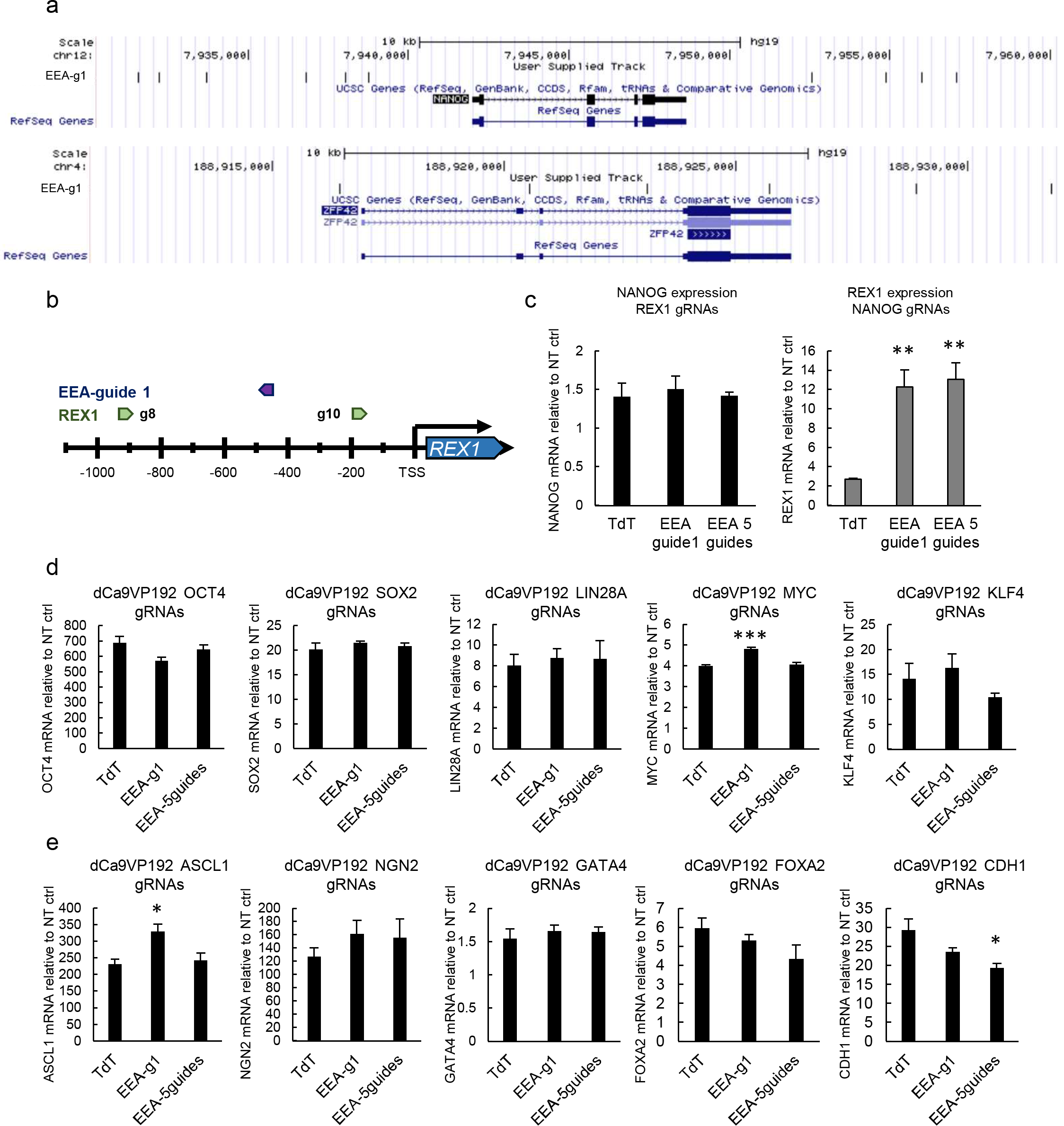
EEA-motif targeting does not improve activation of all pluripotent reprogramming factors. (a) Positions of EEA-gRNA 1 sites near *NANOG* and *REX1* genes. (b) Schematic representation of *REX1* promoter targeting showing *REX1* guide positions and a position of EEA-guide1 between the *REX1* activation guides. (c) Simultaneous targeting of *REX1* and EEA-motif does not affect *NANOG* activation, whereas simultaneous targeting of EEA-motif with *NANOG* gRNAs increases *REX1* activation in an EEA-motif guide dependent manner. (d and e) Simultaneous EEA-motif targeting does not have consistent effect in improving activation of *OCT4*, *SOX2*, *KLF4*, *LIN28A* or *MYC* reprogramming factors (d), or other factors *ASCL1*, *NGN2*, *GATA4*, *FOXA2* or *CDH1* (e). n = 3. Data presented as mean ± SEM, two tailed Student’s t-test. * P<0.05, ** P<0.01, *** P<0.001

**Figure S5.**
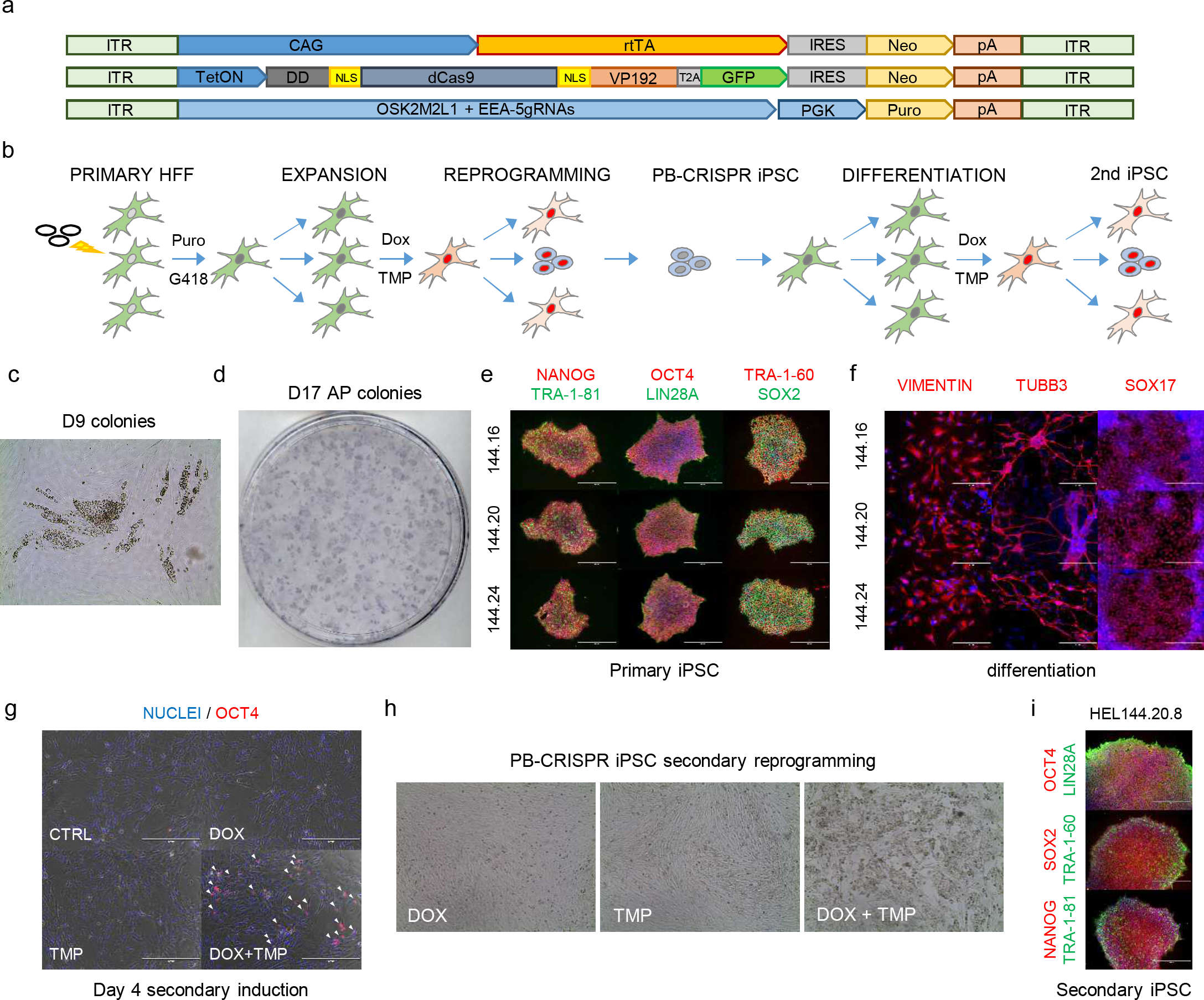
Characterization of PiggyBac CRISPRa iPSC reprogramming. (a) Schematic representation of vectors used in PiggyBac reprogramming. (b) Schematic representation of the reprogramming process. (c) Emerging colonies from primary HFF iPSC induction at day 9 of reprogramming in the presence of doxycycline (DOX) and trimethoprim (TMP). (d) Alkaline phosphatase positive colonies from primary HFF cells at day 17 of reprogramming. (e) Pluripotency marker expression in PB-CRISPRa iPSCs derived from primary HFFs. Scale bar 400μm. (f) Embryoid body differentiation of PB-CRISPRa iPSC into three embryonic germ layer derivatives in vitro. Scale bar 200μm.(g) DOX and TMP dependent activation of CRISPRa targeted OCT4 at day 4 of secondary reprogramming from differentiated fibroblast-like cells derived from HEL144.20. Scale bar 400μm. (h) DOX and TMP dependent morphological changes in fibroblast-like cells differentiated from PB-CRISPR iPSCs. (i) Pluripotency marker expression in secondary PB-CRISPRa iPSCs at passage 3. Scale bar 400μm. Nuclei stained blue.

**Supplemental Table 1.**
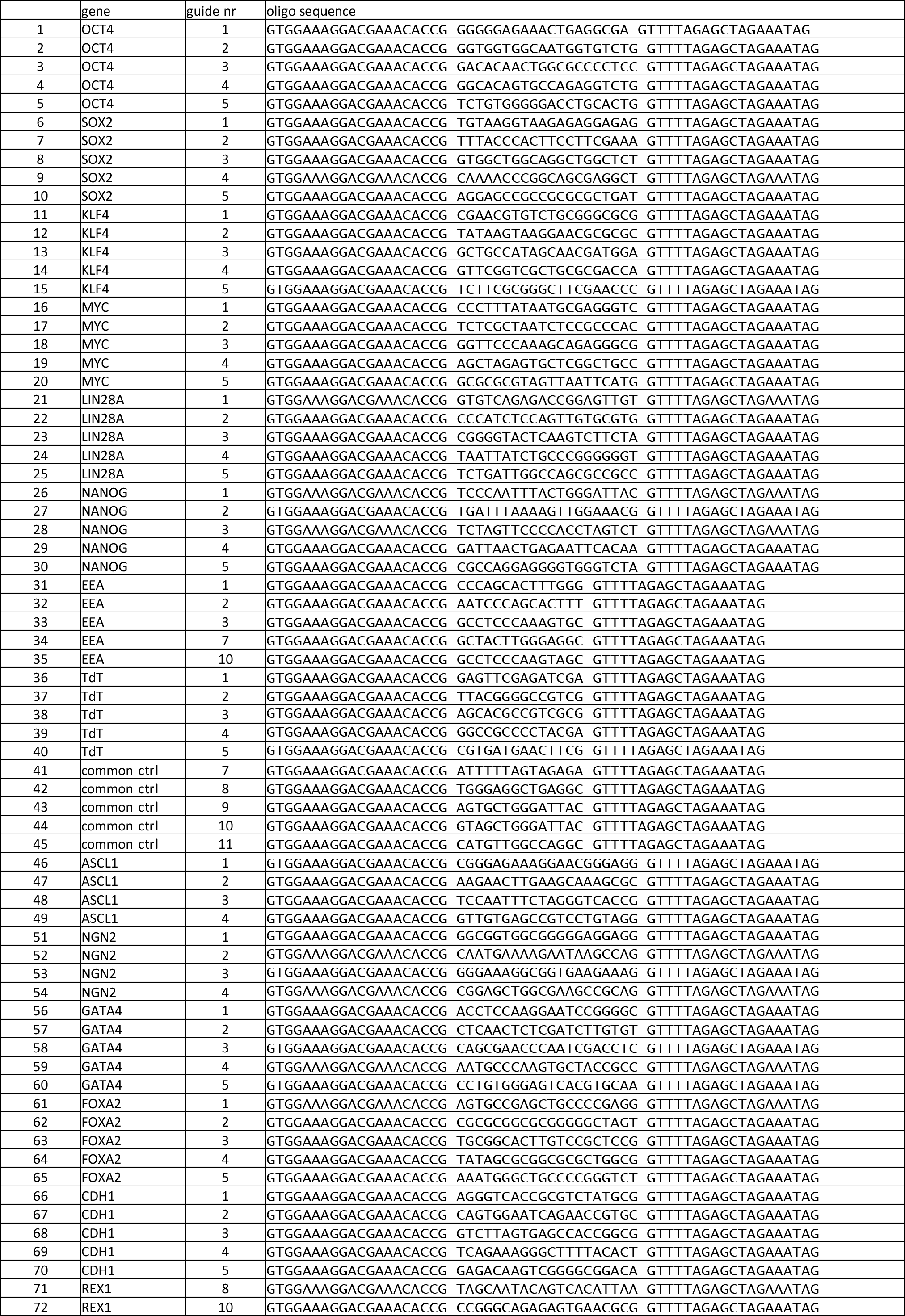
guide RNA oligos

**Supplemental Table 2.** Addgene plasmids

| Addgene ID | Plasmid | Additional info |
| --- | --- | --- |
| 69536 | pCXLE-dCas9VP192-T2A-EGFP | Balboa et al Stem Cell Reports. 2015 Sep 8;5(3):448-59. doi: 10.1016/j.stemcr.2015.08.001. |
| 69535 | pCXLE-dCas9VP192-T2A-EGFP-shP53 | Balboa et al Stem Cell Reports. 2015 Sep 8;5(3):448-59. doi: 10.1016/j.stemcr.2015.08.001. |
| 102885 | PB-CAG-DDdCas9VP192-T2A-GFP-IRES-Neo |  |
| 102886 | PB-CAG-DDdCas9VPH-T2A-GFP-IRES-Neo |  |
| 102887 | PB-CAG-DDdCas9VPP300-T2A-GFP-IRES-Neo |  |
| 102888 | PB-CAG-DDdCas9VPPH-T2A-GFP-IRES-Neo |  |
| 102889 | PB-tight-DDdCas9VP192-T2A-GFP-IRES-Neo |  |
| 102890 | PB-tight-DDdCas9VPH-T2A-GFP-IRES-Neo |  |
| 102891 | PB-tight-DDdCas9VPP300-T2A-GFP-IRES-Neo |  |
| 102892 | PB-tight-DDdCas9VPPH-T2A-GFP-IRES-Neo |  |
| 102893 | PB-GG-OCT4-1-5-PGK-Puro | PiggyBac plasmid. Contains five guides targeting the human OCT4 promoter |
| 102894 | PB-GG-OMKSL-PGK-Puro | PiggyBac plasmid. Contains 3 guides for OCT4, 1 guide for MYC, 1 guide for KLF4, 2 guides for SOX2 and 3 guides for LIN28A promoters. |
| 102895 | pCXLE-dCas9VPH-T2A-GFP-shP53 |  |
| 102896 | pCXLE-dCas9VPP300-T2A-GFP-shP53 |  |
| 102897 | pCXLE-dCas9VPPH-T2A-GFP-shP53 |  |
| 102898 | GG-EBNA-EEA-5guides-PGK-Puro | Replicating episomal plasmid. Contains 5 guides targeting the EEA-motif. |
| 102899 | GG-EBNA-OMKSL-PP | Replicating episomal plasmid. Contains 3 guides for OCT4, 1 guide for MYC, 1 guide for KLF4, 2 guides for SOX2 and 3 guides for LIN28A promoters. Works better when combined with KM plasmid with extra guides for KLF4 and MYC. |
| 102900 | GG-EBNA-KM-PP | Replicating episomal plasmid. Contains 5 guides for MYC and 5 guides for KLF4 promoters. |
| 102901 | GG-EBNA-OS-PP | Replicating episomal plasmid. Contains 5 guides for OCT4 and 5 guides for SOX2 promoters. |
| 102902 | GG-EBNA-OSK2M2L1-PP | Replicating episomal plasmid. Contains 3 guides for OCT4, 2 guides for MYC, 2 guides for KLF4, 2 guides for SOX2 and 1 guides for LIN28A promoters. |
| 102903 | GG-EBNA-TdT-guide1-PGK-Puro | Replicating episomal plasmid. Control guide plasmid. Contains 1 guide targeting the TdTomato sequence. |
| 102904 | GG-EBNA-EEA-guide1-PGK-Puro | Replicating episomal plasmid. Contains guide nr.1 targeting the EEA-motif. |
| 102905 | pCXLE-dCas9KRAB-G-shP53 |  |
| 102906 | pCXLE-dCas9GFP-shP53 |  |
| 102907 | PB-tight-DDdCas9GFP-IRES-Neo |  |
| 102908 | PB-EEA-g1-PGK-Puro | PiggyBac plasmid. Contains guide nr.1 targeting the EEA-motif. |
| 102909 | PB-EEA-5g-OSK2M2L1-PGK-Puro | PiggyBac plasmid. Contains 5 guides targeting the EEA-motif, 3 guides for OCT4, 2 guides for MYC, 2 guides for KLF4, 2 guides for SOX2 and 1 guides for LIN28A promoters. |

**Supplemental Table 3.**
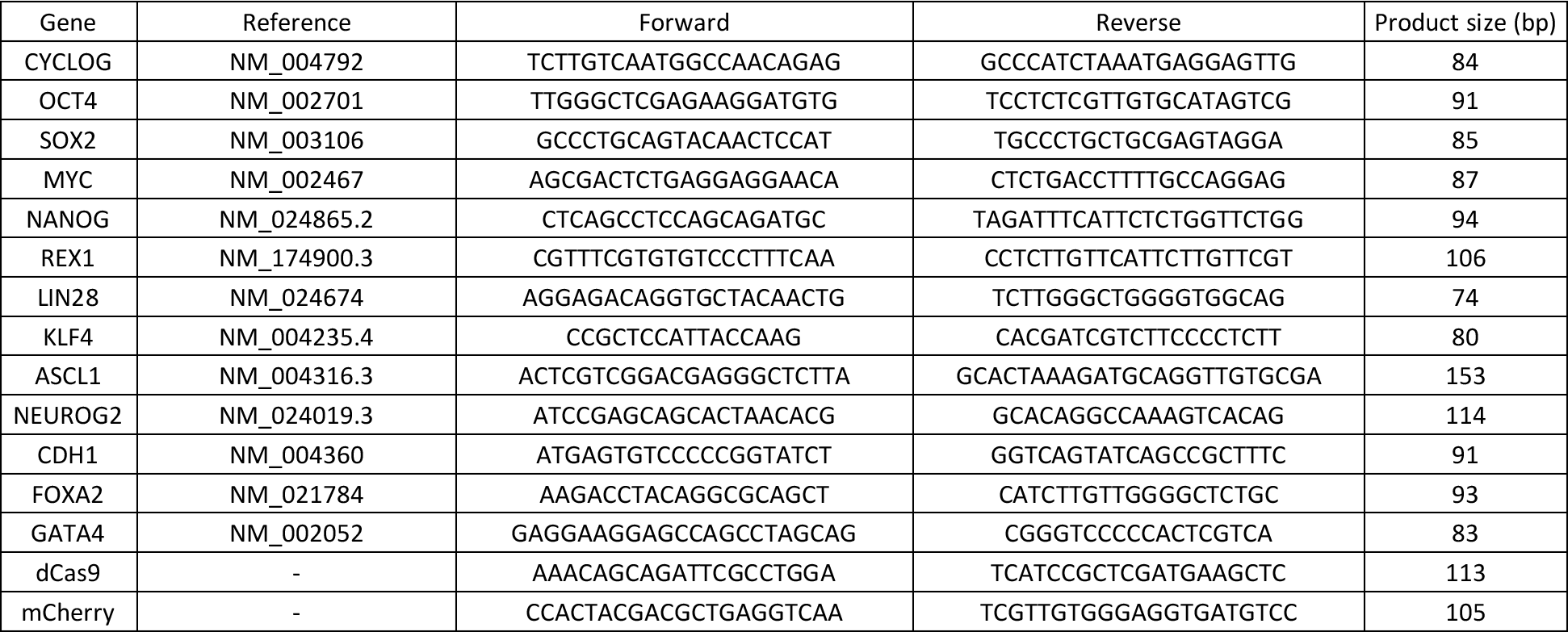
qPCR primers

